# CD73 controls neutrophil responsiveness to type I interferon impairing antibacterial responses during secondary pneumococcal pneumonia

**DOI:** 10.64898/2026.08.12.744509

**Authors:** Alexsandra P. Lenhard, Chloe E. Picciano, Miles J. Stefko, Shaunna R. Simmons, Manmeet Bhalla, Bruce A. Davidson, Elsa N. Bou Ghanem

## Abstract

*Streptococcus pneumoniae* (pneumococcus) are asymptomatic colonizers of the nasopharynx but can progress to pulmonary and systemic pathogens upon influenza A virus (IAV) infection. Polymorphonuclear cells (PMNs) are key for controlling *S. pneumoniae*, but their antibacterial function is impaired by IAV. Using a mouse model that allows transition of pneumococci from colonizers to disease-causing pathogens upon IAV co-infection, we examined the signaling pathways impairing PMN responses. We found that type I interferons (IFN) produced upon IAV infection drives impairment of PMN antibacterial responses as treatment with IFN inhibited pneumococcal killing by PMNs *ex vivo*, and *in vivo* blocking of IFN receptor 1 (IFNAR1) in IAV infected mice rescued PMN function. In exploring what controls PMN responsiveness to IFN, we tested CD73, an ectonucleotidase that regulates PMN responses in primary pneumococcal pneumonia. CD73 was required for the ability of neutrophils to respond to IFN. While wildtype and CD73KO mice had comparable IFNAR expression on PMNs and IFNɑ and IFNꞵ levels upon co-infection, CD73KO PMNs expressed significantly lower levels of the interferon stimulated protein IFIT1 and were less responsive to IFN-mediated inhibition of antimicrobial activity *ex vivo*. In exploring mechanisms, we found that IFN and CD73 elevated reactive oxygen species production by PMN in response to IAV, which impaired their ability to kill pneumococci. Importantly, co-infected CD73KO mice cleared bacteremia and survived significantly better than wildtype controls. These findings suggest that CD73 impairs host defense against IAV/*S. pneumoniae* co-infection in part by sensitizing PMNs to type I IFN-mediated inhibition of antibacterial function.

## Introduction

Secondary bacterial pneumonia increases significantly during flu season [1]. One of the most common bacteria often associated with secondary pneumonia following Influenza A Virus (IAV) infection are *Streptococcus pneumoniae* (pneumococcus) [2,3]. In fact, the risk of developing pneumococcal pneumonia is enhanced 100-fold by IAV infection [4]. *S. pneumoniae* are Gram-positive bacteria that reside in the nasopharynx of most individuals without causing overt disease [2,3]. Within the nasopharynx, *S. pneumoniae* have been reported to form biofilms [5], resulting in asymptomatic colonization of the host. However, upon IAV infection, *S. pneumoniae* can disseminate from the nasopharynx into the lower respiratory tract and establish infection within the lung and systemically [6,7]. Understanding the host factors that play a role in this transition of *S. pneumoniae* from an asymptomatic colonizer to a disease-causing pathogen upon viral infection is crucial for improving infection outcomes.

During primary pneumococcal pneumonia, a hallmark of pulmonary infection is the influx of polymorphonuclear cells (PMNs), also known as neutrophils, to the lungs [8,9]. PMNs are required for control of infection, as depletion of these cells in preclinical models results in higher *S. pneumoniae* burden in the lungs, systemic spread, and detrimental infection outcome [10–12], demonstrating that PMNs are important for control of lung localized infection as well as clearance of disseminated bacteria. These cells are also clinically relevant, as neutropenic patients have a higher risk for developing pneumonia [13]. PMNs respond to *S. pneumoniae* via phagocytosis, production of reactive oxygen species (ROS), degranulation and release of neutrophil extracellular traps (NETs) [14]. We previously found that during IAV/*S. pneumoniae* co-infection, PMNs are recruited to the lungs, but their function is impaired [6]. However, the mechanisms by which viral infection impairs PMN antibacterial responses remains unexplored.

One difference in the host microenvironment during primary versus secondary pneumococcal pneumonia, is the amount of type I Interferons (IFNs) produced. Type I IFNs are produced in response to viral infections and include the cytokines IFNɑ and IFNꞵ, which signal through the heterodimer receptor IFNAR1/2 leading to transcription of interferon stimulated genes (ISGs), which are crucial for anti-viral immune responses and control of viral replication [15–17]. However, type I IFNs may be detrimental for antibacterial host defense, as prior work found that in the absence of IFNAR1, mice are less susceptible to *S. pneumoniae* pulmonary challenge introduced five days following IAV infection [18]. IFNAR1KO mice challenged with IAV had higher recruitment of PMNs to the lungs following *S. pneumoniae* coinfection [18]. However, it is not clear whether type I IFN signaling on PMNs alters their ability to directly kill *S. pneumoniae*. PMNs are known to respond to type I IFN stimulation [19,20] in a tissue and pathogen dependent manner [20,21].

Another key pathway known to control PMN responses during single *S. pneumoniae* and single IAV infection is the Extracellular Adenosine (EAD) pathway [9,12,22–24]. Signaling through this pathway occurs when damaged cells leak ATP, and then two enzymes, CD39 and CD73, de-phosphorylate ATP into adenosine, which can then signal through one of its four G-protein coupled receptors, A1, A2A, A2B, and A3 [25,26]. In primary pneumococcal pneumonia, early adenosine production by CD73 and signaling via A1 receptors was crucial for the ability of PMNs to kill bacteria, and pharmacological inhibition or genetic ablation of CD73 resulted in significantly higher bacterial numbers across host organs and worsened survival upon challenge of mice with *S. pneumoniae* [9,22,27]. In contrast, adenosine production by CD73 and signaling via the A1 receptor following IAV infection did not affect viral loads, but impaired host survival by exacerbating PMN pulmonary influx and lung damage at later timepoints following viral challenge [28,29]. Whether the role of the EAD pathway changes in polymicrobial versus single infections is not known.

The majority of studies examining host responses during *S. pneumoniae/*IAV infection rely on directly delivering the bacteria to the lungs days after IAV infection [30,31]. To better understand the host factors responsible for transition of pneumococcus from asymptomatic colonizer to a disseminating pathogen following viral infection, we established a murine model that allows initial bacterial nasopharyngeal colonization that disseminates upon coinfection with IAV [6,7]. In this model mice are colonized with biofilm-grown *S. pneumoniae* that allows asymptomatic colonization of the nasopharynx, then two days later infected with IAV in both the upper and lower respiratory tract. This model resulted in bacterial spread to the lungs, systemic spread to the blood that was dependent on bacteria strain, and clinical signs of disease and lethality that were dependent on host background and age [7]. Therefore, this model is ideal for parsing out changes in immune responses to polymicrobial infections.

Using this physiologically relevant model, we examined the host signaling pathways controlling PMN responses during IAV/*S. pneumoniae* co-infection. We found that during co-infection, CD73 controls PMN responsiveness to type I IFN signaling, which results in elevated ROS responses that paradoxically impair the ability of PMNs to kill *S. pneumoniae*. We also found that CD73 had no effect on bacterial colonization of the nasopharynx or pathogen load in the lungs, but impaired clearance of bacteremia, suggesting a role in systemic host defense during co-infection.

## Materials and Methods

### Ethics Statement

All animal studies were performed in accordance with the recommendations in the Guide for the Care and Use of Laboratory Animals and the University at Buffalo Institutional Animal Care and Use Committee guidelines (IACUC), approval number MIC33018Y.

### Mice

C57BL/6 wild type (WT) mice (2-3 months) were purchased from Jackson Laboratories. CD73 knockout (CD73KO) mice on a C57BL/6 background [32] were purchased from Jackson laboratories and bred at a specific-pathogen free facility at the University at Buffalo. All the experiments were performed in male mice, as males are more susceptible to *S. pneumoniae* infection [33,34].

### H292 and MDCK Cells

Human pulmonary mucopidermoid carcinoma-derived NCI-H292 (ATCC) “H292” cells were grown at 37°C with 5% CO2 in RPMI 1640 (ATCC) supplemented with 10% FBS (Gibco) and 100U penicillin/streptomycin (ThermoFisher) [35]. H292 cells were fixed prior to use in growth of biofilm by utilizing 4% Paraformaldehyde (ThermoFisher) for 1 hour on ice prior to addition of bacteria [7]. Madin-Darby Canine Kidney (MDCK) cells were purchased from ATCC, and grown in Dulbecco’s Modified Eagle Medium (Corning) supplemented with 100U penicillin/streptomycin (ThermoFisher) and 10% FBS (Gibco) at 37°C with 5% CO2.

### Bacteria

*Streptococcus pneumoniae* serotype 4 (TIGR4 strain) were a kind gift from Andrew Camilli [36]. GFP-expressing *Streptococcus pneumoniae* TIGR4 was a kind gift from Sarah Roggensack [22]. Bacteria were grown in a biofilm as previously described for *in vivo* assays [7], and were dispersed from the biofilm utilizing heat for *ex vivo* assays [37]. Briefly, *S. pneumoniae* were grown in chemically defined media (CDM) at a starting OD of 0.05, to a final OD of 0.2. CDM was prepared as previously described [7]. The bacteria were then added to a 24-well plate of fixed pulmonary H292 epithelial cells and incubated for 48 hours with media changes every 12 hours with fresh CDM media at 34°C/ 5% CO_2_. At 48 hours, the biofilm was harvested, and aliquots were frozen in CDM supplemented with 20% gelatin saved at −80°C until use. Prior to use *in vivo*, bacterial titer was determined. For biofilm-dispersed pneumococcus, at 48 hours of biofilm growth, the plate was placed in an incubator at 38.5°C/5% CO_2_ for 4 hours, the supernatants were then harvested to collect bacteria released from biofilms and aliquots were frozen as described above. Bacterial titers as well as burden in various organs were determined by plating serial dilutions on tryptic soy agar plates supplemented with 5% sheep blood.

### Virus

A mouse-adapted strain of influenza A virus A/PR/8/34 H1N1 was utilized for the experiments [6,38]. To determine viral loads, plaque assays on lung samples were performed utilizing Madin-Darby Canine Kidney (MDCK) cells [7]. Briefly, MDCK cells were added to 6 well plates, allowed to become confluent, then saved lung homogenate supernatants were added to the plates, with each sample being serially diluted. Wells were overlayed with L15 media (Caisson Labs) +1.6% agarose (SeaKem LE, Lonza), plates were placed in an incubator at 37°C with 5% CO_2_ for 48 hours, then stained for plaques utilizing crystal violet as previously described [7].

### Infection and Treatments

Un-anesthetized mice were intranasally colonized with 10^7^ biofilm grown pneumococcus in a 10uL volume delivered directly to the nares (5ul/nares). Two days following colonization, mice were anesthetized using isoflurane, then intratracheally infected with 20 PFU/50uL IAV using the tongue-pull method resulting in oropharyngeal aspiration as previously described [7]. Following recovery from anesthesia, mice were intranasally infected with 200 PFU/10uL IAV. Two days following IAV infection, mice were euthanized and organs harvested to determine bacterial and viral burden and for immune cell analysis via flow cytometry [7]. Another group of mice were monitored for clinical symptoms of disease including respiratory quality, weight, activity, grooming, posture, and breathing [9,39] up to 10 days post viral infection. Where indicated, at day 0 and +1 with respect to viral infection, mice were treated intraperitoneally (i.p.) with ɑ-IFNAR1 antibody (MAR1-5A3, BioXcell) or isotype control (IgG1, _K_, BioXcell) at a concentration of 250ug/100uL.

### PMN Isolation

PMNs were isolated from the bone marrow of naïve or IAV infected mice two days following infection. In brief, the femurs and tibias of the mice were flushed with RPMI supplemented with 10% FBS and 2mM EDTA, followed by lysis of the red blood cells and then remaining cells were washed and resuspended in PBS. Density centrifugation was utilized to isolate the PMNs via histopaque 1119 (Sigma) and histopaque 1077 (Sigma) as described previously [22,40]. Following isolation, PMNs were resuspended to a desired concentration of 2.5×10^6^/mL in Hank’s Balanced Salt Solution with no calcium or magnesium supplemented with 0.1% gelatin.

### Opsonophagocytic Killing Assay

PMN antimicrobial activity was measured using an *ex vivo* opsonophagocytic killing assay as previously described [23,41]. Briefly, PMNs were incubated with biofilm-dispersed bacteria at a multiplicity of infection (MOI) of 0.01 in the presence of 3% sera from matching mice, rotating at 37°C for 40 minutes. Where noted, prior to incubation with the bacteria, PMNs were treated with recombinant IFNɑ and IFNꞵ (R&D Systems^TM^), at varying doses noted on the figures or Diphenyleneiodonium chloride (DPI) (Sigma) at 10*_μ_*M for 35 minutes rotating at 37°C. Reactions were plated on tryptic soy agar plates supplemented with 5% sheep blood to determine bacterial numbers. Percent bacterial killing was calculated with respect to no PMN controls under the same treatment conditions.

### IFN ELISAs

The lungs from the various mouse groups were collected in PBS, homogenized, and spun down to collect the supernatants. Type I IFNɑ and IFNꞵ levels in the supernatants were measured utilizing commercially available kits (R&D Systems^TM^) as per manufacturer’s instructions. Absorbance (450nm) was read on a Biotek Plate reader.

### *In vitro* and *In vivo* Reactive Oxygen Species (ROS) Assays

Total ROS was measured *in vitro* using a chemiluminescent plate based assay as previously described [27]. In brief, following PMN isolation, cells were resuspended in Hank’s Balanced Salt Solution with no Ca^2+^ or Mg^2+^ and allowed to rest for 1 hour at room temperature. PMNs were then spun down and resuspended in KRP buffer (Phosphate buffered saline with 5mM glucose, 1mM CaCl_2_ and 1mM MgSO_4_) and allowed to rest for an additional 30 minutes. Following the rest, PMNs were added to a 96 well white LUMITRAC plates (USA Scientific). PMNs were either treated with 3% sera alone (uninfected), or 3% matching sera with biofilm-dispersed *S. pneumoniae*. Following infection, 50μM or Isoluminol or Luminol plus 10U/ml HRP (Sigma) were added to the wells to detect extracellular and intracellular ROS respectively. Luminescence was immediately measured in a Biotek plate reader for 1 hour at 37°C.

Mitochondrial ROS was measured *in vitro* using MitoSox (Invitrogen) assay as previously described [23] with some modifications. Isolated PMNs were rested in KRP buffer for 30 minutes as described above. Following rest, PMNs were added to a non-tissue culture (TC) treated round-bottom 96 well plate. PMNs were then incubated with 5μM MitoSOX-Red fluorescent dye (Invitrogen) for 15 minutes at 37°C and then either treated with 3% sera alone (uninfected), or 3% matching sera with biofilm-dispersed *S. pneumoniae*. Fluorescence was immediately measured in a Biotek plate reader for 1 hour at 37°C.

ROS *in vivo* was determined via flow cytometry utilizing the fluorescent dye CellROX Deep Red (Invitrogen C10422). Briefly, following organ processing, samples from the bone marrow and lungs were stained with Fc Block (2.4G2, Invitrogen), CellROX Deep Red, and Ly6G (1A8, Biolegend) for 20 minutes at 37°C. Following the incubation, samples were washed and resuspended in FACS buffer, filtered, and immediately ran on the BD FACS LSRFortessa^TM^ and at least 25,000 events were analyzed using FlowJo.

### *In vitro* Phagocytosis Assay

To determine phagocytosis, PMNs isolated from the bone marrow were infected with biofilm-dispersed GFP-*S. pneumoniae*, or biofilm-dispersed wild type *S. pneumoniae* as a negative control, for 10 minutes rotating at 37°C. The cells were fixed with BD Cytofix^TM^ buffer on ice. To differentiate associated versus engulfed bacteria, the cells were stained with anti-pneumococcal serotype 4 capsular antibodies raised in rabbit (Cederlane) then with PE-conjugated secondary anti-Rabbit IgG antibody (Invitrogen). Following the incubation, samples were washed and resuspended in FACS buffer and analyzed using a BD FACSCelesta^TM^ and FlowJo software to determine the percent of PMNs that had bacteria associated (all GFP+ PMNs) versus engulfed (GFP+, PE-).

### Isolation of Cells for Flow Cytometry

Mice were perfused with 10ml PBS, the lungs removed, washed in PBS, and minced into small pieces. The lungs were then digested for 1 hour with RPMI 1640 supplemented with 10% FBS, 1 mg/ml Type II collagenase (Worthington), and 50 U/ml Deoxyribonuclease I (Worthington) at 37° C/ 5% CO_2_. Single-cell suspensions were obtained by mashing the digested lungs. Total bone marrow cells were collected by flushing the femurs and tibias with 10ml RPMI 1640 supplemented with 10% FBS and 2mM EDTA. The red blood cells were removed by treatment with a hypotonic lysis buffer (Lonza). Cells were analyzed using flow cytometry.

### Flow Cytometry

Bone marrow, blood, and lung single cell suspensions were incubated with Fc Block (Clone 2.4G2) and Live/Dead dye (Invitrogen), then stained with the following antibodies: CD45 (30-F11, BD Bioscience), Ly6G (1A8, Biolegend), IFNAR1 (MAR1-5A3, Invitrogen), IFNAR2 (FAB1083P, R&D), IFIT1 (OTI3G8, Novus Biologicals) and CD73 (eBIO TY/11.8, BD Bioscience). Following staining, cells were fixed with BD Cytofix^TM^ buffer on ice. Fluorescent intensities were measured on the BD FACS LSRFortessa^TM^ and at least 25,000 events were analyzed using FlowJo. For PMN effector activity, following Fc Block and L/D stain, cells were surface stained for Ly6G (1A8, Biolegend) and CD63 (NVG-2, Biolegend) then treated with BD Cytofix/Cytoperm^TM^ buffer on ice. Permeabilized cells were then stained with antibodies against Neutrophil Elastase (JF098-6, Novus Biologicals) or Citrullinated H3 (NB10057135, Novus Biologicals) followed by PE-conjugated secondary anti-Donkey IgG antibody (Invitrogen). Percentage and fluorescence intensities were measured on a Cytek Aurora and at least 25,000 events were analyzed using FlowJo.

### Statistics

All statistical analyses were performed using GraphPad Prism. Viral and bacterial burden were log transformed. Normality and Lognormality tests were performed on data using D’Agostino-Pearson and Shapiro-Wilk analyses. For data normally distributed, significant differences were determined with one-way ANOVA followed by Šídák’s or Dunnett’s multiple comparisons test, or unpaired t test where applicable. For data not normally distributed, significant differences were determined using Kruskal-Wallis followed by Dunn’s multiple comparisons test where applicable. Incidence of bacteremia was determined using Fisher’s exact test. Survival analysis was performed using a Logrank (Mantel-Cox) test. For fold changes, One-Sample t test was used to test significant differences from one. *p* values < 0.05 were deemed to be significant. * denote *p* <0.05 ** *p* <.01, *** *p* <.001 and **** *p* <.0001.

## Results

### PMNs are responsive to type I IFNs produced during IAV/*S. pneumoniae* co-infection

As type I IFNs are a crucial part of the antiviral response [42], we wanted to determine if they were produced during IAV/pneumococcal co-infection. We measured the levels of IFNɑ and IFNꞵ in the sera and lungs of uninfected, singly colonized (+Sp), single viral (+IAV), and IAV/*S. pneumoniae* co-infected wild type C57BL/6 (WT) mice at 48 hours post co-infection, as indicated in the experimental scheme (Figure 1A). We found that IFNɑ levels in the lungs and circulation of WT mice increased with single viral infection and co-infection compared to baseline (Figure 1B and C). IFNꞵ levels within the lungs and circulation of infected WT mice were comparable to uninfected controls (Figure 1D and E). Next, we wanted to assess if PMNs respond to type I IFN produced during infection. We first measured the expression of Interferon Alpha and Beta Receptor Subunit 1 and 2 (IFNAR1 and IFNAR2) on WT PMNs (gating strategy in Supplemental Figure 1). As during infection, PMNs must egress from the bone marrow to the site of infection [43,44], and to get a comprehensive picture of PMN responses, we assessed bone marrow, blood, and pulmonary PMNs. We found that the percentage of bone marrow WT PMNs expressing IFNAR was very low (Figure 2A-D). In contrast, we found that as WT PMNs egress from the bone marrow and enter circulation, they upregulate IFNAR receptors (Figure 2E-H). The percentage of PMNs expressing IFNAR1 upon co-infection increased from 10% in the bone marrow to 40 and 60% in the circulation and lungs respectively (Fig 2A, E and I). IFNAR2 positive PMNs, increased from <1% in the bone marrow to around 5 and 15% in the circulation and lungs (Fig 2C, G and K). The amount of IFNAR1 and 2 expressed, as measured by geometric mean florescence intensity (gMFI), also significantly increased in co-infected mice upon entry to the circulation and lungs (Fig 2B, F, J and Fig. 3). Infection did not affect the percentage of WT PMNs expressing these receptors nor the amounts of receptors expressed on PMNs (Figure 2) in any organ. These findings suggest that upon exit from the bone marrow, PMNs upregulate expression of IFNAR 1 and 2.

**Figure 1.**
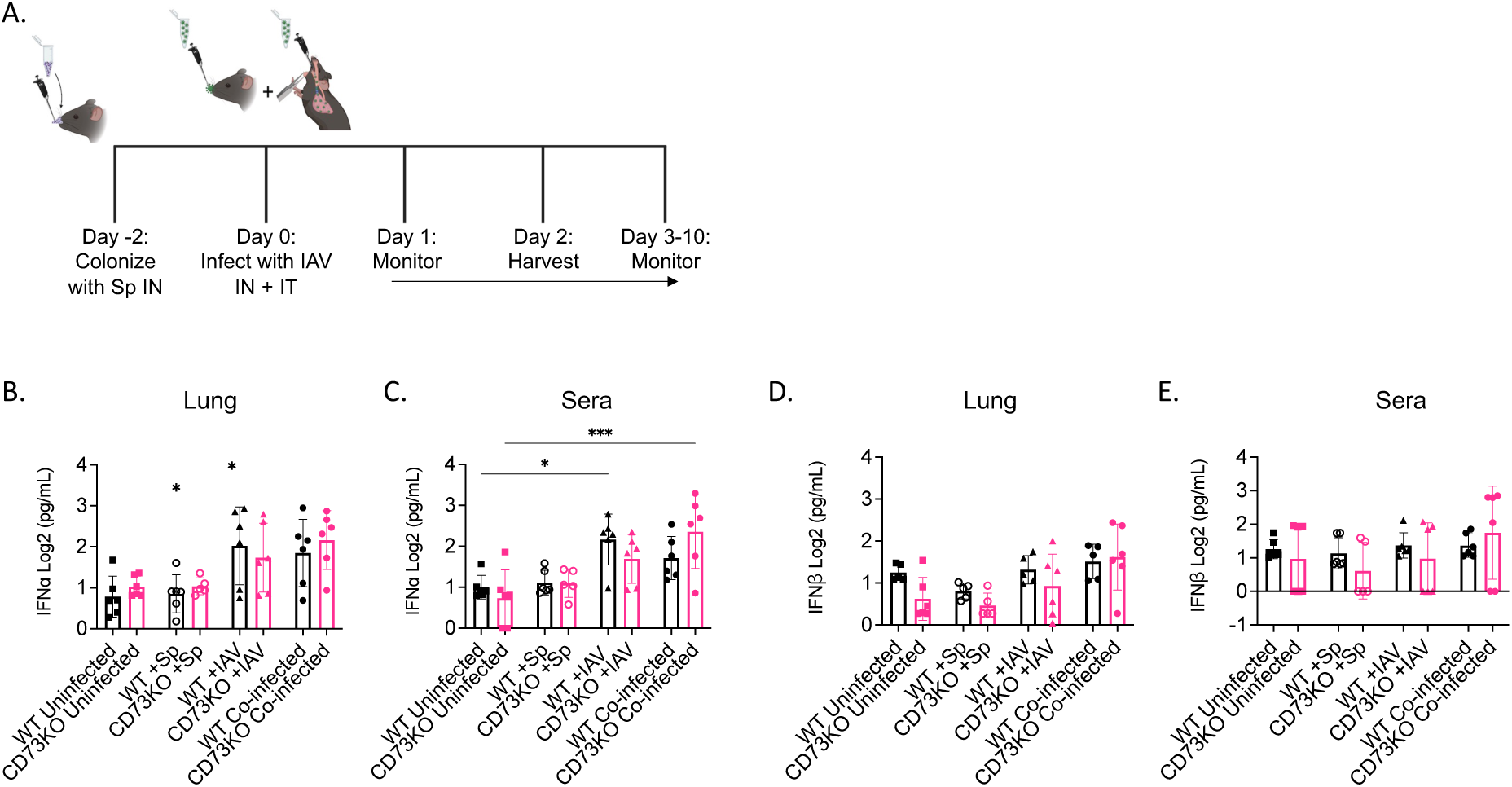
IAV/*S. pneumoniae* co-infection elicits production of type I IFNs. C57BL/6 (WT) or CD73KO mice were colonized with *S. pneumoniae* and two days later infected with influenza A Virus (co-infected) following the timeline shown in (A). Control groups included mice that were infected with virus alone (+IAV), colonized with *S. pneumoniae* alone (+Sp), or mock-infected with PBS (uninfected). Two days post IAV infection, levels of IFNɑ (B, C) and IFNꞵ (D, E) in the lungs and circulation were measured by ELISA. Data are pooled from two separate experiments with n=6 mice per group. *Denotes significant differences between indicated groups as determined by One-way ANOVA followed by Šídák’s multiple comparisons test.

**Figure 2.**
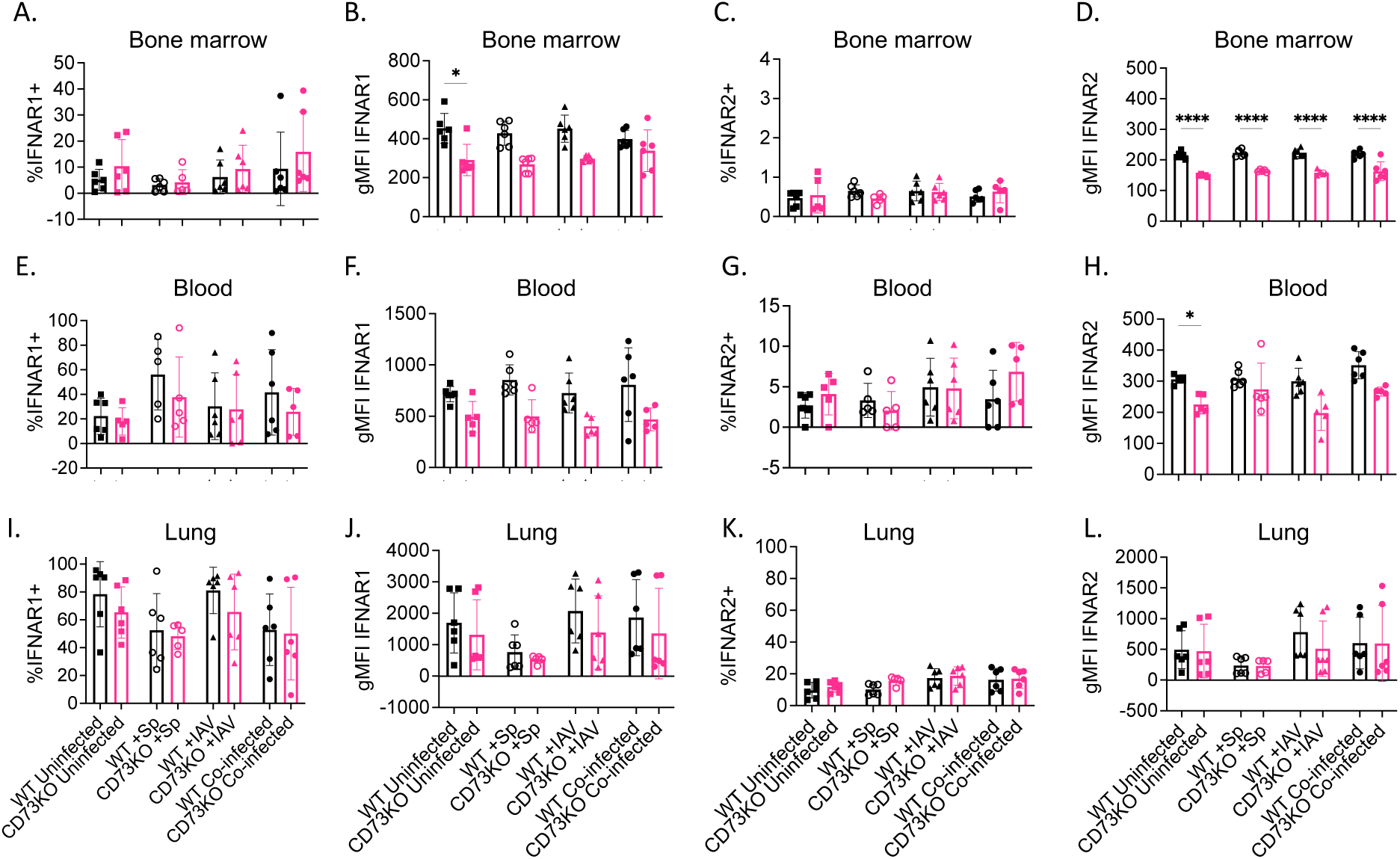
Bone marrow PMNs from CD73KO mice express lower levels of IFNAR compared to wild type controls. WT or CD73KO mice were either co-infected with *S. pneumoniae* and Influenza A Virus (co-infected), infected with virus alone (+IAV), colonized with *S. pneumoniae* alone (+Sp), or mock-infected with PBS (uninfected). Two days post IAV infection, the bone marrow (A-D), blood (E-H) and lungs (I-L) were harvested and assessed by flow cytometry for percent % and expression (gMFI) of IFNAR1 and IFNAR2 on PMNs. Data are pooled from two separate experiments with n=5-6 mice per group. *Denotes significant differences between indicated groups as determined by One-way ANOVA followed by Šídák’s multiple comparisons test.

**Figure 3.**
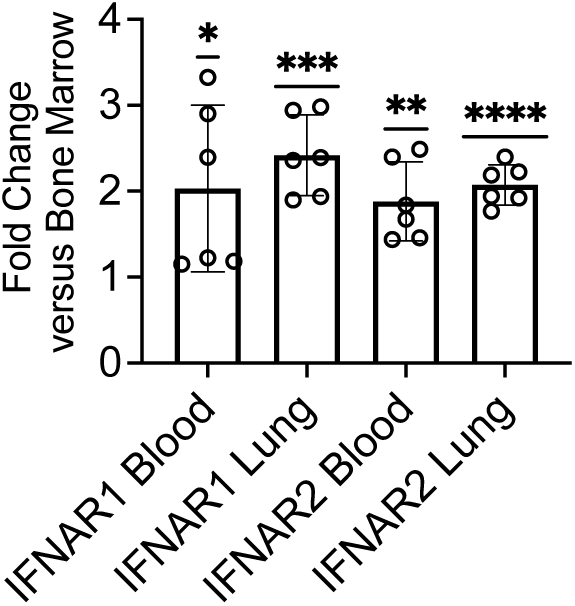
PMNs upregulate IFNAR expression following exit from the bone marrow. Fold change of IFNAR1 and IFNAR2 on PMNs in the blood and lung of co-infected WT mice were determined relative to their expression level in the bone marrow under the same challenge conditions. Data are pooled from two separate experiments with n=6 mice per group. *Denotes significant differences in fold changes as determined by One-Sample t test.

To investigate PMN responsiveness to type I IFNs, we then looked at the percentage (Figure 4A-C) and expression (Figure 4D-F) of IFIT1, an interferon stimulated gene (ISG) reported to be upregulated on PMNs upon treatment with type I IFNs [45]. We found that the percentage of WT PMNs expressing IFIT1 in the bone marrow (Figure 4A), blood (Figure 4B), and lungs (Figure 4C) significantly increased in response to co-infection compared to uninfected controls. Overall, these findings suggest that WT PMNs respond to type I IFNs produced during IAV/*S. pneumoniae* co-infection.

**Figure 4.**
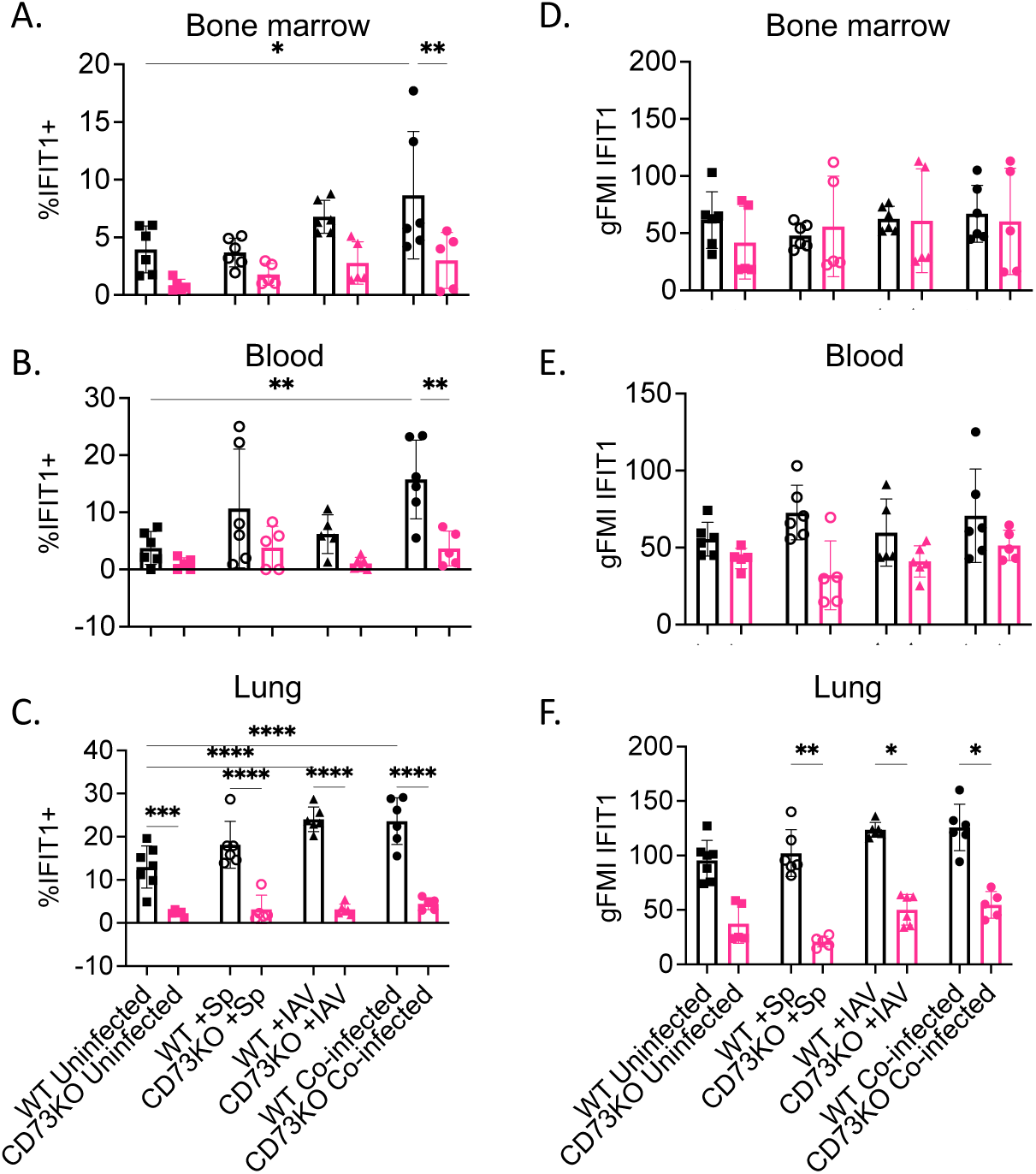
PMNs from CD73KO mice express lower levels of IFIT1 compared to wild type controls. WT or CD73KO mice were either co-infected with *S. pneumoniae* and Influenza A Virus (co-infected), infected with virus alone (+IAV), colonized with *S. pneumoniae* alone (+Sp), or mock-infected with PBS (uninfected). Two days post IAV infection, the bone marrow, blood, and lungs were harvested and assessed by flow cytometry for percent % (A-C) and expression (gMFI) (D-F) of IFIT1 on PMNs. Data are pooled from two separate experiments with n=5-6 mice per group. *Denotes significant differences between indicated groups as determined by One-way ANOVA followed by Šídák’s multiple comparisons test.

### PMN antibacterial killing is impaired by type I IFNs

Previous work found that PMNs from an IAV infected host are significantly impaired in their ability to kill *S. pneumoniae* [6]. We wanted to investigate if impairment in PMN function in an IAV infected host is driven by type I IFNs. To do this, we performed an opsonophagocytic killing assay where PMNs from naïve WT mice were treated with varying doses of recombinant IFNɑ and IFNꞵ, reflective of what we found *in vivo* during infection (Figure 1A-B). We used biofilm-dispersed bacteria to model *in vivo* conditions, as dispersion results in upregulation of bacterial virulence factors [46], that allow *S. pneumoniae* to establish pulmonary and invasive infections. We found that addition of IFNɑ or IFNꞵ or both, impaired PMN antibacterial killing (Figure 5A). We next tested if blocking type I IFN signaling *in vivo*, can we rescue PMN function. To address this, IAV infected mice were treated with an anti-IFNAR1 blocking antibody at the time of viral infection, and one day post infection, and the ability of PMNs to kill *S. pneumoniae ex vivo* was measured [9]. We found that as previously reported [6], IAV infection completely abrogated the ability of PMNs to kill *S. pneumoniae* (Figure 5B). However, when IFNAR1 was blocked, PMN anti-bacterial activity was restored to levels comparable to those of PMNs isolated from naïve hosts (Figure 5B). Overall, these findings indicate that in an IAV infected host, PMN anti-bacterial function is impaired by type I IFNs.

**Figure 5.**
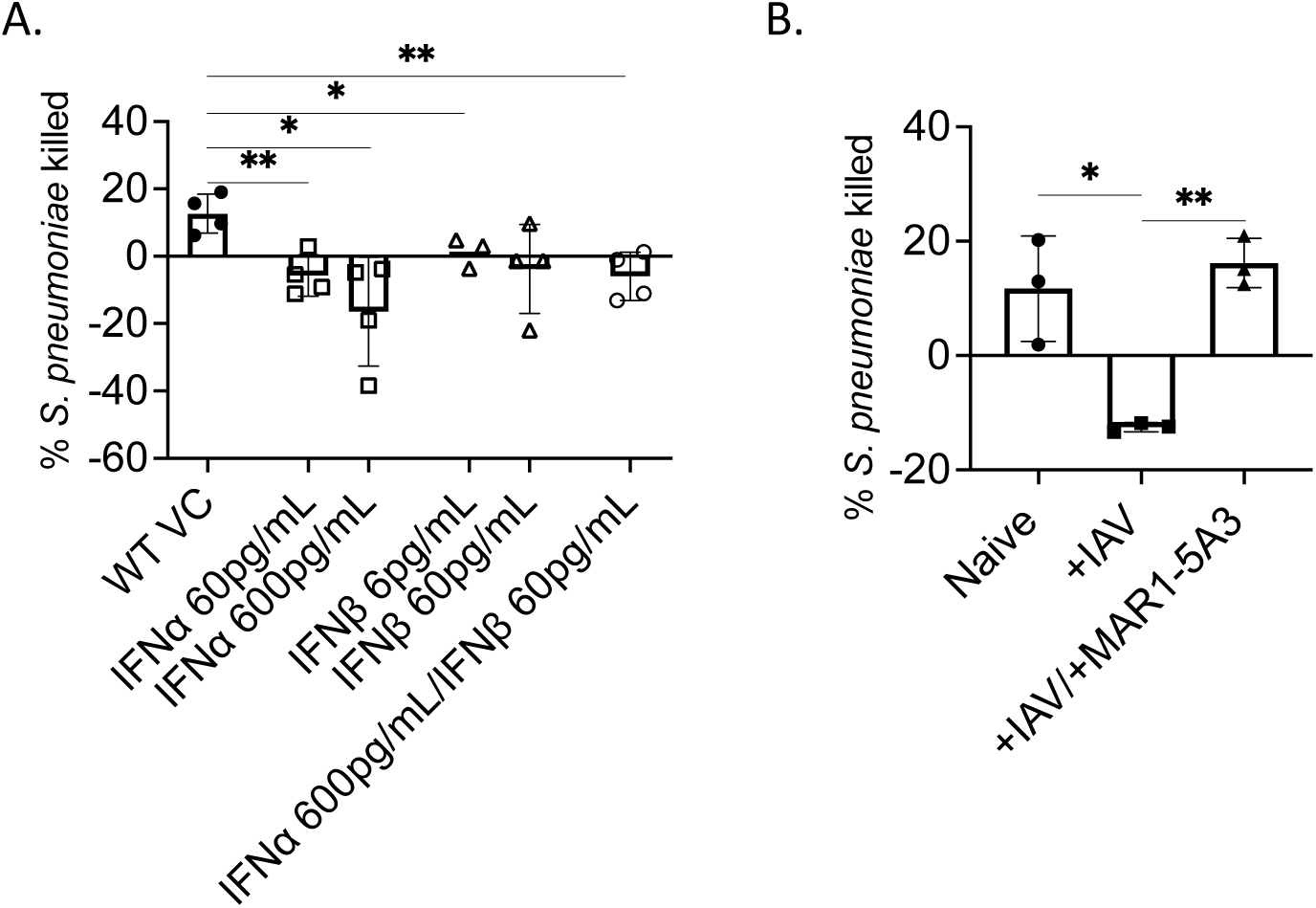
Type I IFNs impair the ability of PMNs to kill *S. pneumoniae*. PMNs isolated from the bone marrow of WT mice were mock-treated (VC) or treated with exogenous IFNɑ at 60 or 600 pg/mL, or IFNꞵ at the 6 or 60 pg/mL, or a combination of IFNɑ at 600pg/mL and IFNꞵ at 60pg/mL (A). Following treatment, PMNs were infected with biofilm-dispersed *S. pneumoniae* and the percentage of bacteria killed was calculated in comparison to a no PMN control for each condition. WT mice were mock infected (naïve) or IAV infected mice and treated with isotype controls or IFNAR1 blocking antibody (MAR1-5A3) at days 0, and +1 with respect to infection (B). Two days post IAV challenge, PMNs were isolated from the bone marrow and challenged with biofilm-dispersed *S. pneumoniae.* Percentage of bacteria killed was calculated in comparison to a no PMN control for each condition. Data are pooled from four (A) and three (B) separate experiments and each dot represents the average of technical replicates from an individual mouse. *Denotes significant differences between indicated groups as determined by One-way ANOVA followed by Dunnett’s multiple comparisons test.

### PMNs express CD73 during IAV/*S. pneumoniae* co-infection

We next wanted to test what controls PMN responsiveness to type I IFNs. We had previously found that CD73 controls PMN responses during primary pneumococcal pneumonia [12,22,23,26,35,47]. However, if that was also true during IAV/pneumococcal co-infection had not been tested. To address this, we first looked at CD73 expression on PMNs during co-infection (gating strategy in Supplemental Figure 1). We found that 40% of PMNs in the bone marrow expressed CD73, and that CD73 expression was significantly downregulated on PMNs in response to co-infection (Figure 6A, B), which recapitulated previous findings during primary pneumococcal pneumonia [27]. Within circulation, the percentage of PMNs expressing CD73 did not change in response to infection (Figure 6C, D). In the lungs, the majority of PMNs expressed CD73 on their surface, with no significant change based on infection (Figure 6E, F). Overall, these findings show that during IAV/pneumococcal co-infection, CD73 is consistently expressed on the surface of circulating and pulmonary PMNs.

**Figure 6.**
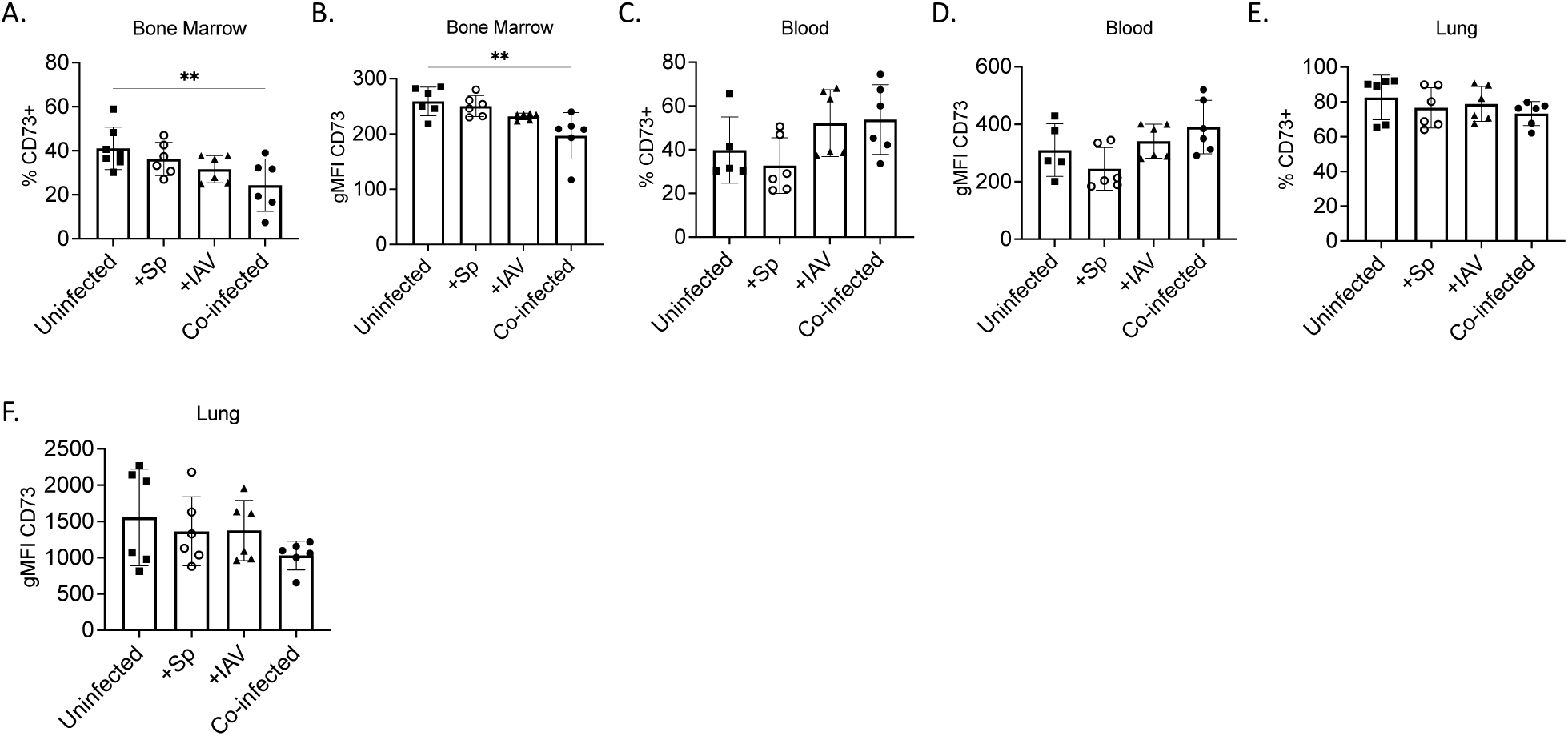
PMNs from uninfected and co-infected hosts express CD73. WT mice were either co-infected with *S. pneumoniae* and Influenza A Virus (co-infected), infected with virus alone (+IAV), colonized with *S. pneumoniae* alone (+Sp), or mock-infected with PBS (uninfected). Two days post IAV infection, the bone marrow (A, B), blood (C, D) and lungs (E, F) were harvested and assessed by flow cytometry for percent % and expression (gMFI) of CD73 on PMNs. Data are pooled from two separate experiments with n=6 mice per group (A-F). *Denotes significant differences as determined by One-way ANOVA followed by Šídák’s multiple comparisons test.

### CD73 controls sensitivity to type I IFNs on PMNs

We next wanted to investigate if there was an association between CD73 and PMN responses to type I IFN. To address this, we compared type I IFN levels and receptor expression on PMNs from WT versus CD73KO mice. We found that there was no significant difference in IFNɑ or IFNꞵ levels within the lungs across all conditions between the mouse strains (Figure 1B-E). When we examined PMNs, we found there was no difference in the percentages or numbers of PMNs in the bone marrow and lungs of WT and CD73KO mice (Supplemental Figure 2). When we investigated IFNAR1 and IFNAR2 receptor expression, we found that there was no difference in the percentage of PMNs that express these receptors between WT and CD73KO mice (Figure 2A, C, E, G, I and K). The expression of both receptors was significantly lower on bone marrow CD73KO PMNs compared to WT controls (Figure 2B, D), however these differences were no longer apparent in the circulation (Figure 2F, H) or lungs (Figure 2J, L). We did, however, find that across all organs, during co-infection, CD73KO mice had a significantly lower percentage of IFIT1+ PMNs (Figure 4A-C), as well as lower levels of IFIT1 in circulating and pulmonary PMNs compared to WT controls (Figure 4D-F). These data suggest that in the absence of CD73, PMNs are less responsive to type I IFNs. To address this functionally, we performed an opsonophagocytic killing assay with treatment of recombinant type I IFNs comparing PMNs isolated from WT or CD73KO mice, using cytokine levels found during *in vivo* infection (Fig. 1B and C). We found that while type I IFN treatment impaired the antibacterial activity of WT PMNs, CD73KO PMNs were less sensitive to treatment and were able to kill *S. pneumoniae* even when stimulated with type I IFNs (Figure 7). These findings suggest that CD73 drives sensitivity to type I IFN on PMNs.

**Figure 7.**
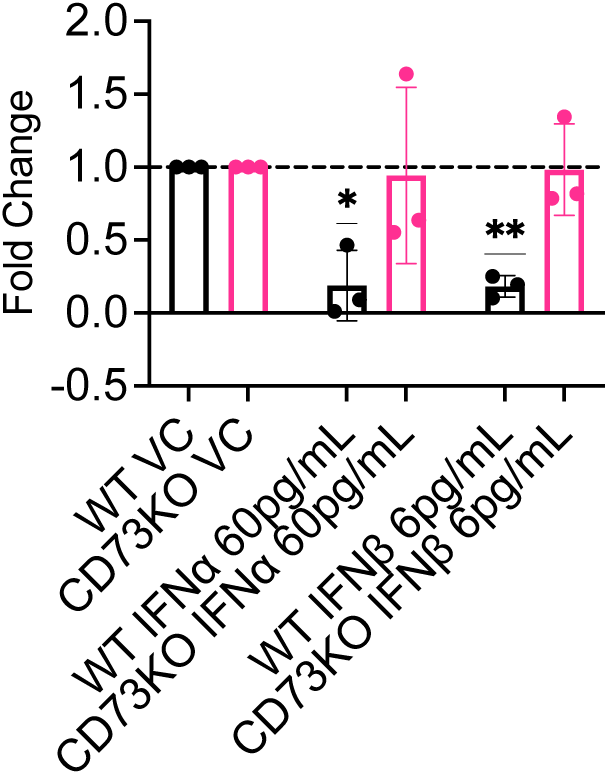
CD73KO PMNs are less responsive to type I IFN-mediated impairment of antibacterial activity. PMNs were isolated from the bone marrow of WT or CD73KO mice and treated with exogenous IFNɑ at 60 or 600 pg/mL or IFNꞵ at the 6 or 60 pg/mL or mock-treated (VC). Following treatment, PMNs were infected with biofilm-dispersed *S. pneumoniae* and the percentage of bacteria killed was calculated in comparison to a no PMN control for each condition. Fold changes in bacterial killing in response to IFN treatments were then calculated with respect to VC condition for each mouse. Data are pooled from three separate experiments and each dot represents the average of technical replicates from an individual mouse. *\** Denotes significant differences in fold changes as determined by One-Sample t test.

### IAV infection increases intracellular ROS production in PMNs

PMNs kill pathogens through many different effector functions. These include phagocytosis, production of reactive oxygen species (ROS), and release of antimicrobials from granules and neutrophil extracellular traps (NETs) [14]. As PMNs from IAV infected WT hosts have impaired ability to kill *S. pneumoniae*, we wanted to investigate which effector function(s) are impaired. To address this, we first measured uptake of the bacteria from bone marrow WT PMNs isolated from a naïve or IAV infected host *ex vivo*. Utilizing GFP-expressing bacteria and inside out staining followed by flow cytometry as we described previously [22], we found no difference in the ability of WT PMNs isolated from a naïve or IAV infected host to associate with (Figure 8A) or engulf (Figure 8B) *S. pneumoniae*.

**Figure 8.**
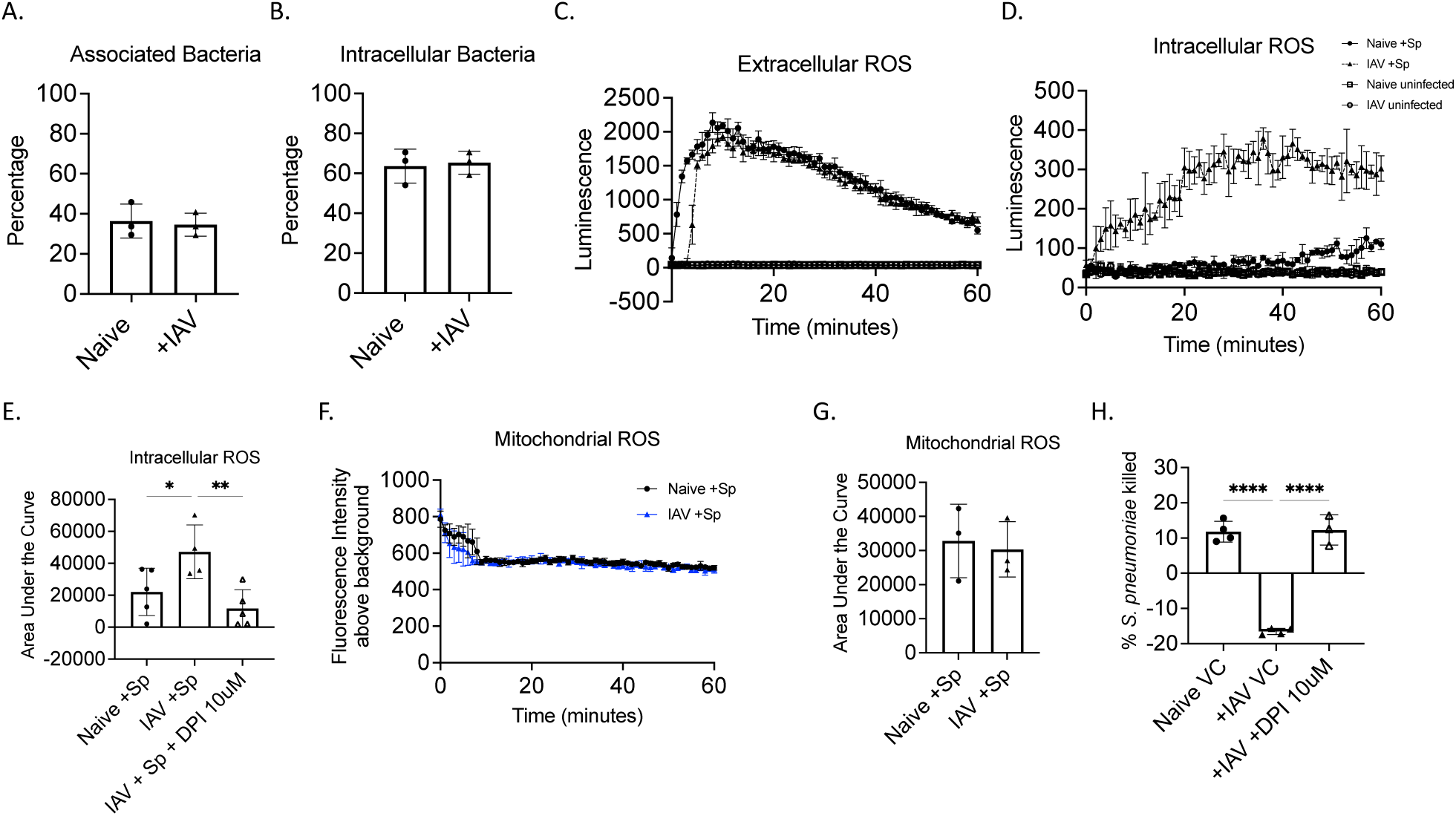
IAV infection increases ROS production and impairs bacterial killing by PMNs. PMNs were isolated from naïve and IAV infected WT mice two days post infection and then challanged with biofilm-dispersed GFP-*S. pneumoniae* (A-B). Following challenge, PMNs were analyzed by flow cytometry to determine the percentage of PMNs with associated (A) versus engulfed (B) bacteria using inside out staining with anti-*S. pneumoniae* capsular antibodies. PMNs were isolated from naïve and IAV infected WT mice two days post infection (C-G). Cells were then mock-treated or challenged with biofilm-dispersed *S. pneumoniae* (+Sp) and total extracellular (C) as well as total intracellular (D-E) ROS production determined by isoluminol and luminol staining respectively, while mitochondrial (F-G) ROS production determined by MitoSox. For mitochondrial ROS, background production by uninfected controls are subtracted from the data. Representative data are shown from one of three separate experiments (C, D and F). Area under the curve was calculated to determine differences between PMNs infected with biofilm-dispersed pneumococcus from naïve or IAV infected hosts and data were pooled from three separate experiments (E, G). PMNs were isolated from naïve and IAV infected C57BL/6 mice two days post infection and treated with VC or 10uM of DPI (a NADPH oxidase inhibitor) for 20 minutes prior to challenge with biofilm-dispersed *S. pneumoniae* (H). The percentage of bacteria killed was then calculated in comparison to a no PMN control for each condition. Data are pooled from four separate experiments and each dot represents the average of technical replicates from an individual mouse (H). *Denotes significant differences as determined by One-way ANOVA followed by Šídák’s multiple comparisons test.

We next investigated if IAV infection altered ROS responses in PMNs by using chemiluminescent based assays [23,27]. We found that upon stimulation with *S. pneumoniae*, WT PMNs from an IAV infected host had similar levels of extracellular ROS production (Figure 8C) but had significantly elevated levels of intracellular ROS compared to PMNs from a naïve host (Figure 8D, E). Prior work from our group found that the source of ROS source is imperative for proper control of pneumococcus by PMNs, and that while mitochondrial ROS enhances bacterial killing by PMNs, ROS produced via the NADPH Oxidase impairs the ability of PMNs to kill *S. pneumoniae* [23]. We compared mitochondrial ROS production by WT PMNs from an IAV infected or naïve host in response to *S. pneumoniae* stimulation and found no difference in production (Figure 8F, G), suggesting that the elevated intracellular ROS production in response to IAV is NADPH Oxidase-derived. Indeed, when WT PMNs from IAV infected mice were treated with the NADPH oxidase inhibitor Diphenyleneiodonium Chloride (DPI), their ability to produce intracellular ROS in response to *S. pneumoniae* stimulation was reduced to naïve host levels (Figure 8E). To test whether NADPH Oxidase-derived ROS production was impairing the antibacterial activity of PMNs from IAV infected hosts, we performed an opsonophagocytic killing assay where PMNs were treated with DPI, then incubated with biofilm-dispersed *S. pneumoniae*. We found that DPI treatment rescued the ability of WT PMNs from an IAV infected host to kill *S. pneumoniae* (Figure 8H). Overall, these findings suggest that the IAV-driven impairment in WT PMN function is caused by dysregulation of NADPH-derived ROS production.

To confirm our findings *in vivo*, so we measured ROS production in PMNs via flow cytometry utilizing the fluorescent dye CellROX (gating strategy in Supplemental Figure 3). We found that WT PMNs in the bone marrow begin to produce ROS in response to co-infection (Figure 9A), but that circulating (Figure 9B) and pulmonary (Figure 9C, D) PMNs significantly increased ROS production in response to single IAV infection and co-infection. Overall, these finding demonstrate that IAV infection also elicits ROS production by WT PMNs *in vivo*.

**Figure 9.**
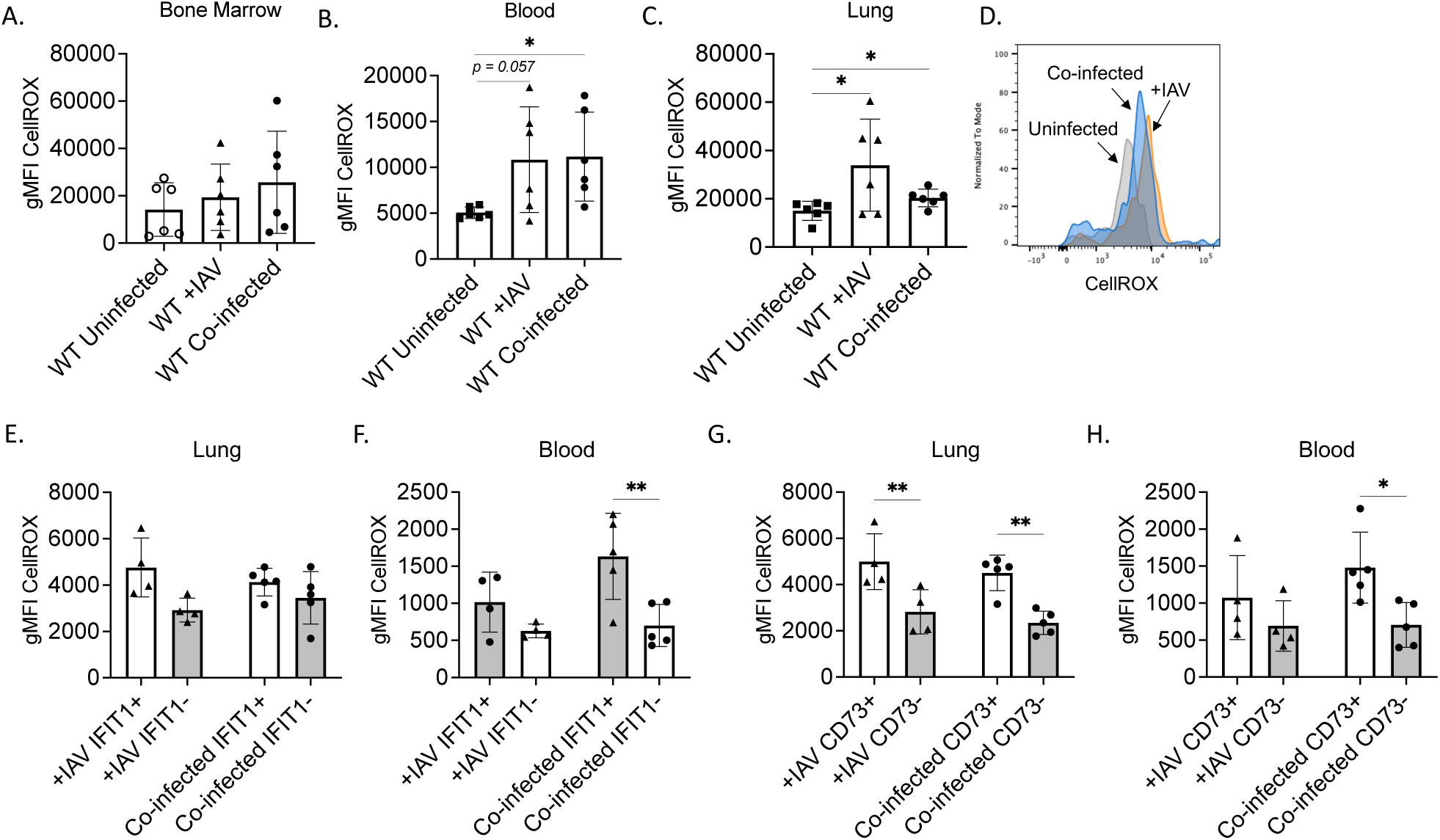
CD73 expression on PMNs enhances ROS production in response to single IAV and IAV/pneumococcal co-infection. WT mice were either co-infected with *S. pneumoniae* and Influenza A Virus (co-infected), infected with virus alone (+IAV), or mock-infected with PBS (uninfected). Two days post IAV infection, the bone marrow, blood, and lungs were harvested and assessed by flow cytometry for expression of CD73 and IFIT on PMNs as well as production of reactive oxygen species (ROS) using CellROX by PMNs. The gMFI of CellROX in total (A-D), IFIT positive versus negative (E-F) and CD73 positive versus negative (G-H) PMNs was determined. Data are pooled from two separate experiments with n=6 mice (A-D) or 4-5 mice (E-H) per group. *Denotes significant differences as determined by One-way ANOVA frollowed by Šídák’s multiple comparisons test.

### IFIT+ PMNs in the circulation produce more ROS during IAV/ *S. pneumoniae* co-infection

We then investigated if the increased ROS production by PMNs in WT co-infected hosts was driven by type I IFNs. We examined ROS produced by PMNs that expressed IFIT1 versus those that did not (gating strategy in Supplemental Figure 3). We found that there was a slight, but not significant increase in the levels of ROS (gMFI CellROX) in IFIT1+ compared to IFIT1-PMNs (Figure 9E) in the lungs. However, we did find that there was a significant increase in levels of ROS (gMFI CellROX) in IFIT1+ compared to IFIT1-PMNs from the blood (Figure 9F). These results suggest that increased ROS during IAV/pneumococcal co-infection in WT mice is driven by type I IFN responsiveness on PMNs in circulation, not the lungs.

### CD73 controls ROS production during IAV/ *S. pneumoniae* co-infection

As we previously found that CD73 controls ROS production by PMNs during primary pneumococcal infection [27], we wanted to investigate if the increased intracellular ROS observed during co-infection was also regulated by CD73. To address this, we measured total ROS produced by CD73+ versus CD73-PMNs via flow cytometry (gating strategy in Supplemental Figure 3). We found that in both IAV and co-infected WT hosts, CD73+ PMNs produced significantly more ROS compared to CD73-PMNs in the lungs and circulation (Figure 9G, 8H). To confirm this, we then compared ROS production in response to infection in WT versus CD73KO mice. We found that while the majority of WT PMNs from the bone marrow and lungs upregulated ROS production in response to infection, CD73KO PMNs failed to do so (Figure 10A, C). Within the circulation, we found that CD73KO PMNs upregulated ROS production in response to IAV single infection, but not co-infection as seen in WT PMNs (Figure 10B). We also compared other PMN effector activities in WT versus CD73KO hosts and found that CD73 had no significant effect on the levels of neutrophil elastase or CD63 (as a proxy for release of primary granules) in response to co-infection (Supplemental Figures 4-6). In line with its effect on ROS production, CD73 was required for the elevated percentage of citrullinated H3 (as a proxy for NETosis) expressing PMNs in the lungs, circulation and bone marrow in response to IAV and IAV/*S. pneumoniae* co-infection (Supplemental Figures 4C, 5C and 6C). Overall, these findings suggest that CD73 is driving the increased ROS production by PMNs in response to IAV/*S. pneumoniae* co-infection.

**Figure 10.**
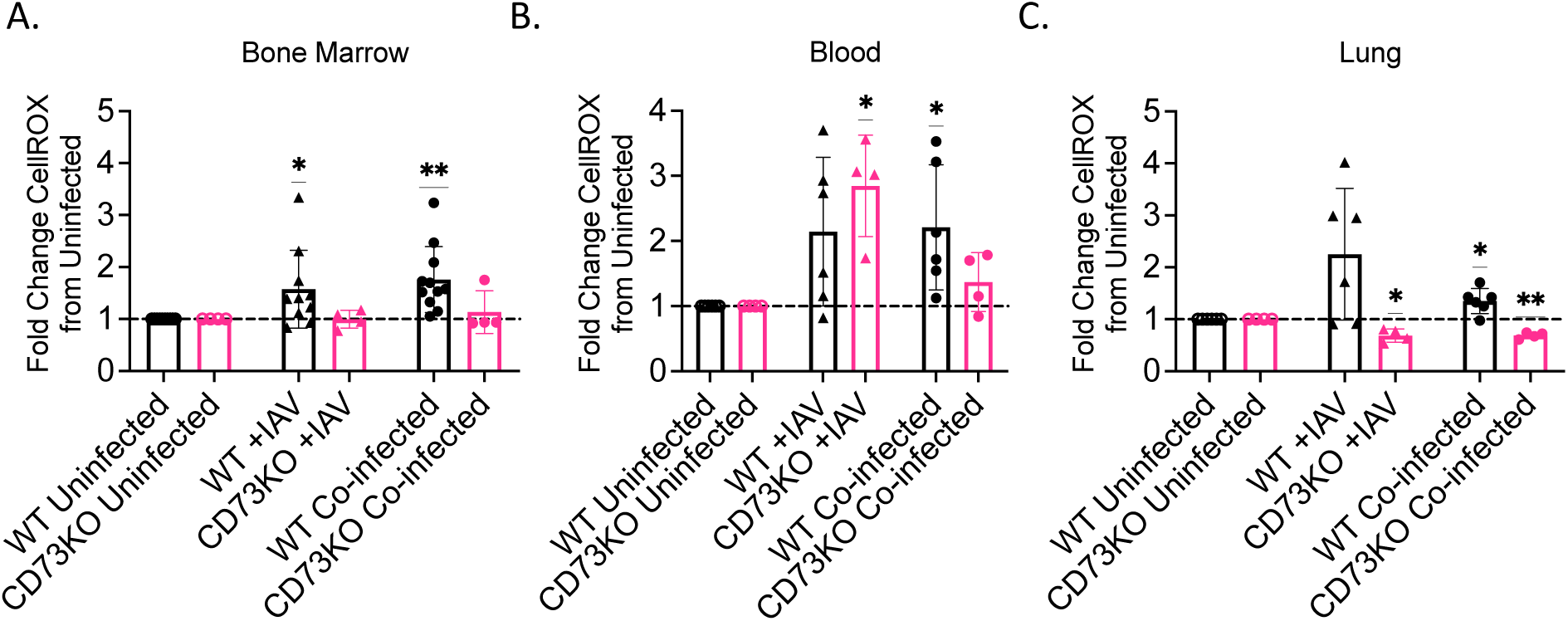
PMNs from CD73KO co-infected mice do not increase ROS production in response to infection. WT and CD73KO mice were either co-infected with *S. pneumoniae* and Influenza A Virus (co-infected), infected with virus alone (+IAV), or mock-infected with PBS (uninfected). Two days post IAV infection, the lungs, bone marrow, and blood were harvested and assessed by flow cytometry for ROS production (CellROX) by PMNs (A-C). Fold change of CellROX gMFI on PMNs in the indicated organs was determined relative to uninfected controls following the indicated challenge conditions for each mouse strain. Data are pooled from two separate experiments with n=4-6 mice per group (A-C). *Denotes significant differences as determined by One-Sample t test.

### CD73 is detrimental to the overall outcome of IAV/ *S. pneumoniae* co-infection

Finally, we tested if CD73 contributes to host susceptibility to IAV/pneumococcal infection. We compared pathogen loads and host symptoms in CD73KO and WT mice in response to IAV/*S. pneumoniae* co-infection. We found that at 48 hours post viral infection, there was no difference in bacteria bound to the nasal tissue or bacteria present within the nasal wash between CD73KO and WT mice in both single colonization and co-infection (Figure 11A). Similarly, the incidence of bacterial spread to the lungs in both WT and CD73KO co-infected mice did not vary (Figure 11B). We also found no difference in the pulmonary viral load in either single IAV or IAV/*S. pneumoniae* co-infection between CD73KO and WT mice (Figure 11C). However, when we measured bacterial dissemination to the systemic circulation, we found that while initial incidence of bacteremia at 48 hpi did not differ, when WT co-infected mice became bacteremic, they succumbed to the infection, whereas in the CD73KO mice were able to control bacterial numbers and clear the infection in the blood (Figure 11D). When we tracked clinical symptoms to determine probability of survival, we found that CD73KO mice survived IAV/*S. pneumoniae* co-infection significantly better than WT mice. We observed that while the majority of WT mice succumbed to the co-infection, more than half of CD73KO mice survived the challenge (Figure 11E). Overall, these findings indicate that CD73 is detrimental to host survival during IAV/*S. pneumoniae* co-infection and plays a role in systemic, rather than respiratory tract host defense against bacteria during co-infection.

**Figure 11.**
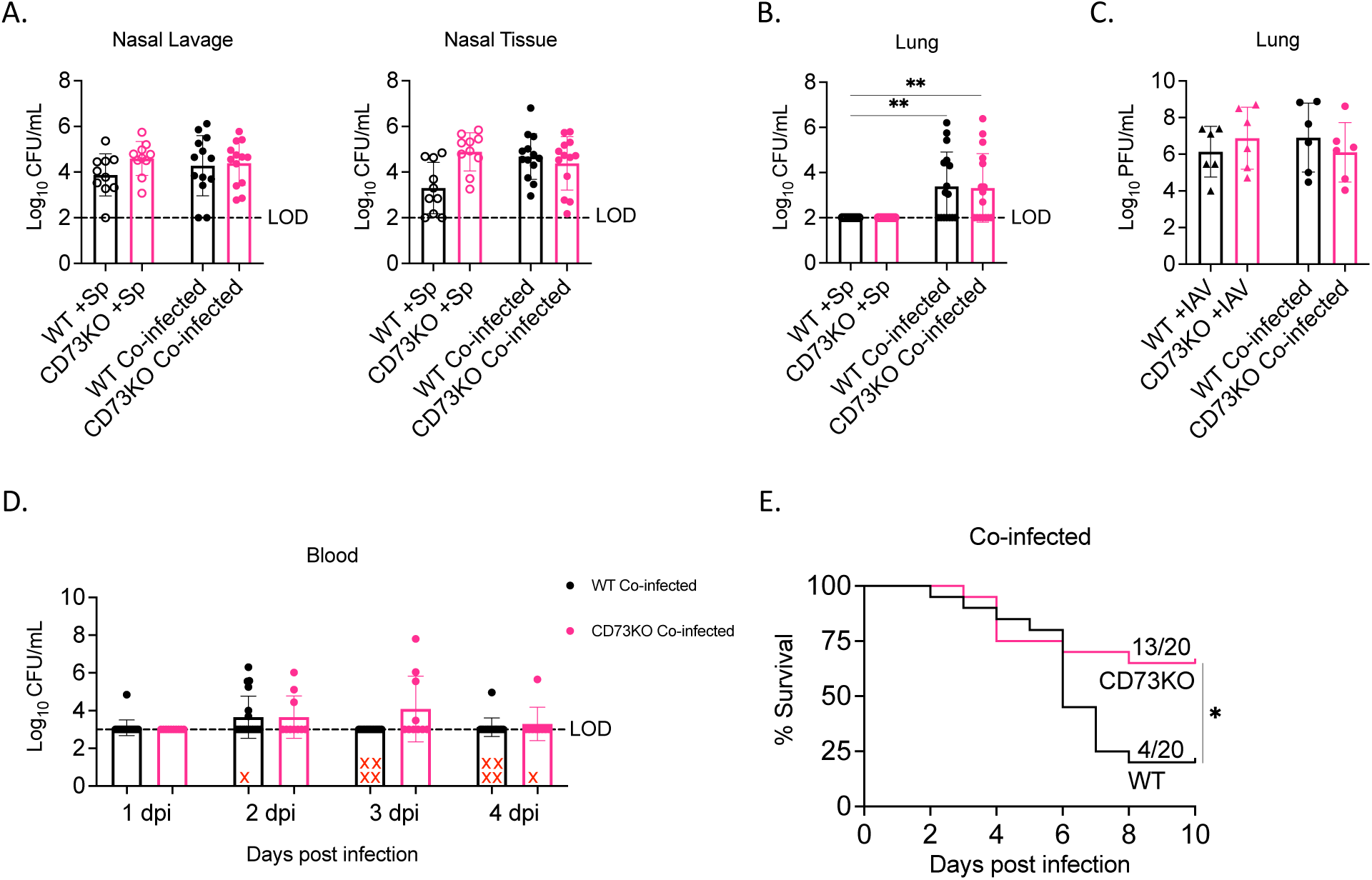
CD73 is detrimental to host survival following IAV/pneumococcal co-infection. WT and CD73KO mice were intranasally colonized with biofilm-grown *S. pneumoniae* TIGR4 or PBS, and then 2 days later were infected with IAV or PBS intranasally and intratracheally. A cohort of mice were euthanized and harvested two days post viral infection to determine bacterial burden in the nasopharynx (A) as well as bacterial (B) and viral (C) burden in the lungs. Bacteremia was measured over time (D). Mice were monitored in co-infected groups for survival for up to 10 days post infection (E). Data are pooled from three separate experiments with n=10 mice for bacterial enumeration (A-B), n=5 mice for viral enumeration (C) and n= 20 mice for tracking bacteremia and survival (D, E). *Denotes significance as determined by Kruskal-Wallis test followed by Dunn’s multiple comparisons test (B) and Log rank (Mantel-Cox) test (F).

## Discussion

Secondary bacterial pneumonia is often a sequalae following influenza A virus infection, with *S. pneumoniae* being a major causative agent during past flu pandemics [2,3,5]. The infection model used in this study mimics what occurs in people, where prior bacterial colonization is considered a pre-requisite for invasive disease [48], and pneumococcus residing asymptomatically in the nasopharynx can transition to a disease-causing pathogen upon viral infection [6,7]. Here we used this model to identify signaling pathways that control neutrophilic responses and host outcome following co-infection. We identified the CD73/ type I IFN axis as drivers of host susceptibility to IAV/pneumococcal co-infection. CD73 did not affect pathogen loads in the upper and lower respiratory tract but rather impaired systemic responses and reduced bacterial clearance from the circulation. We report for the first time that CD73 controls PMN responsiveness to type I IFNs produced upon viral infection which results in impaired bacterial killing paradoxically driven by increased NADPH oxidase-derived ROS. This provides novel insights into the complex signaling networks controlling PMN responses during polymicrobial infections.

PMNs are essential for clearance of *S. pneumoniae* during single bacterial infection [8,9,12,22,23,27,49] and one of the mechanisms by which influenza A infection increases the risk of pneumococcal pneumonia is through impairment of PMN antibacterial function [6,50,51]. In prior work, we found that PMN depletion had no effect on host survival following IAV/pneumococcal co-infection because their anti-bacterial function was impaired [6]. Here we identified type I IFNs, which are required for control of viral replication [15–17], as drivers of this impairment. Similar to prior reports [50,52], we found that IAV infection elicits production of IFNɑ in the lungs and circulation and that a subset of PMNs were responsive to type I IFNs. It is now appreciated that PMNs are heterogenous, and several different subtypes have been identified [45,53]. In human IAV infection, PMN subsets with upregulated interferon-related gene signatures can be indicators for severe disease [54–56]. A subset of PMNs that increase expression of ISGs in response to *Escherichia coli* infection, denoted as G5b PMNs, has also been identified in both humans and mice [45]. Similarly, here, we found that 20% of circulating and pulmonary PMNs upregulated expression of IFIT1, an ISG, in response to *S. pneumoniae*/ IAV coinfection. Importantly, we found that blocking IFNAR1 restored the PMN antibacterial activity in IAV infected hosts. This is in line with recent reports that type I IFN signaling impairs PMN anti-pneumococcal activity in the middle ear during viral infection [50]. Further, several studies have found that while type I IFN is required for pulmonary barrier integrity and host defense against primary pneumococcal pneumonia [57,58], its role is flipped during secondary pneumococcal pneumonia, where IFNAR signaling impairs bacterial clearance during IAV [18,52,59,60] and other respiratory viral infections [61].

We found here that PMN responsiveness to type I IFN is dependent on CD73 expression. CD73 is an ectonucleotidase that is required for conversion of ATP into adenosine in the extracellular space. Very few prior studies have explored the connection between CD73 and type I IFNs. Both IFNɑ and IFNꞵ have been reported to increase expression of CD73 on non-immune endothelial cells, enhancing barrier integrity during inflammation [62–64]. In non-infectious, acute lung injury models that result in hypoxia, IFNɑ was reported to increase CD73 expression on endothelial cells and in the lungs and this was shown to protect against tissue damage [63]. IFNꞵ also resulted in upregulation of CD73 on cultured lung samples, and in patients with acute respiratory distress, intravenous administration of this factor significantly decreased acute mortality [65]. Here we found the opposite, where CD73 was required for the ability of type I IFNs to blunt PMN antibacterial activity, which impaired systemic clearance of the bacteria following co-infection, rendering CD73 detrimental for host survival. We are also the first to report the reverse connection, where CD73 controls responsiveness to type I IFNs in the context of IAV/*S. pneumoniae* co-infection. The mechanisms by which CD73 does that is unknown and may be via transcriptional control of IFIT1, as we previously found that CD73KO PMNs have dysregulated transcriptional responses to *S. pneumoniae* challenge [47] and prior work found that CD73 was required for transcription of IFNɑ at mucosal sites [62].

In exploring the mechanisms of PMN impairment, we found that reactive oxygen species (ROS) production by PMNs is increased, and that this is detrimental to bacterial clearance by these cells during co-infection with IAV. The role of ROS, specifically NADPH oxidase-derived ROS, in viral infections has been found to be beneficial early on for control of viral replication [66]. However, we previously reported that in primary pneumococcal pneumonia, NADPH oxidase-derived ROS impairs the ability of PMNs isolated from healthy donors to kill pneumococci, while mitochondrial-derived ROS is crucial for control of these bacteria [23]. Here we found that IAV infection did not alter mitochondrial ROS production, but significantly increased NADPH oxidase-derived ROS, and that inhibition of the NADPH oxidase reversed IAV-driven impairment of PMN antibacterial function. Therefore, we identified a PMN effector activity that while is beneficial for control of virus, impairs host defense against bacteria in the context of polymicrobial infections. The enhanced ROS production observed here was driven by CD73 and responsiveness to type I IFNs. We previously found that CD73 was needed for production of intracellular ROS by PMNs in response to single *S. pneumoniae* infection [27]. Similarly, it has also been shown that addition of IFNɑ primes ROS production by PMNs in response to several stimuli [21]. Therefore, it is possible that CD73 mediates ROS production via control of PMN responsiveness to IFNɑ, as we found increased ROS during *in vivo* co-infection in PMNs that were CD73 and IFIT positive. The mechanisms by which this is occurring and whether that is via increase expression or assembly of the NADPH oxidase complex components is yet to be determined.

The role of CD73 in host outcome is altered during single versus polymicrobial infection. In the context of single IAV infection, CD73 was reported to have an immunomodulatory effect by altering chemotaxis of immune cells [28]. Once ATP is leaked from damaged cells it is sequentially de-phosphorylated into extracellular adenosine by CD39 and CD73 [26]. EAD can then act on one of its four G-protein coupled receptors that have varying affinity to this ligand [26]. During single IAV infection, signaling through the high affinity A1 receptor was detrimental to host survival, contributing to increased PMN recruitment to the lung and causing severe lung pathology [29]. Our lab previously found that in contrast, CD73 and signaling through the A1 receptor is beneficial for early host defense against primary pneumococcal pneumonia [9,35,47]. Adenosine production by CD73 regulated pulmonary PMN influx into the lungs during primary pneumococcal pneumonia and was required for the ability of these cells to kill and clear the bacteria [9,35,47]. In contrast, we found that overt accumulation of EAD [12], and signaling through the low affinity A2B receptor later on during pneumococcal infection, impairs PMN function and host survival [23]. Here, during IAV/pneumococcal co-infection, we found that CD73 impairs host survival by reducing bacterial clearance from the circulation, without affecting PMN pulmonary recruitment. Therefore, it is possible that in the presence of IAV, CD73 results in high levels of EAD that accumulates early on during infection, activating the low affinity A2B receptor, thus impairing antibacterial defense.

There are limitations to the work presented here. One limitation is our focus on a single bacterial and viral strain. There are many serotypes of *S. pneumoniae*, including more than 100 identified to date [67] and their prevalence and disease potential in humans varies [67]. In prior work, we did test several strains of pneumococcus across various serotypes and found that strain type is important in disease severity during co-infection, as the more invasive strains, such as

TIGR4 used in this study, were the ones to show bacterial dissemination to the lungs and blood in viral infected hosts [7]. Therefore, we focused here on TIGR4 as it allowed us to examine local pulmonary as well as systemic immune responses. Similarly, there are many strains of IAV that differ in their pathogenicity [68–71] and here we focused on the mouse adapted H1N1 PR8 virus [7]. Therefore, it would be relevant to study how viral strain alters host susceptibility to secondary pneumococcal pneumonia moving forward. Biological sex should also be considered as a limitation of this study, as this can greatly affect contractibility and outcome to various diseases [72–74]. In this work we used male mice due as they are the more susceptible sex. In the context of both primary and secondary pneumococcal pneumonia, males are more susceptible to infection [34,75], therefore our focus on male hosts is scientifically justified. In future studies, it would be important to investigate if biological sex alters the signaling pathways involved in IAV/pneumococcal co-infection outcome. Finally, another limitation here is the focus on an animal model. While use of mice is beneficial for understanding host response to infections, and this model mimic the kinetics of pneumococcal colonization and subsequent transition to disease causing pathogen upon IAV infection that we observe in people [7], there are differences in the immune response between mice and humans [76]. Using experimental human colonization models, studies found that immune responses are altered during co-infections and that nasal exposure to a live-attenuated influenza vaccine strain impaired PMN anti-pneumococcal activity [51,77], similar to what we report here, validating the relevance of this model. Therefore, the work presented here is relevant as it phenocopies human findings and can be a tool for finding and testing therapeutics to improve outcome of disease. In summary, this work can have implications for interventions for polymicrobial infections, as we identified two signaling pathways elicited by IAV infection, namely type I IFN and the extracellular adenosine producing CD73, that impair antibacterial host defense and that can be targeted pharmacologically [26].

## Acknowledgements

We would like to acknowledge Andrew Camilli, John Leong and Sara Roggensack for bacterial strains.

## Author contributions

APL conducted research, analyzed data and wrote paper. CEP, MJS, SRS and MB conducted research and analyzed data. BAD provided valuable reagents and feedback. ENBG designed research, wrote the paper and had responsibility for final content. All authors read and approved the final manuscript.

## Data Availability

All data are available from the corresponding author upon request.

