## Supplemental Data for "CD73 controls neutrophil responsiveness to type I interferon impairing antibacterial responses during secondary pneumococcal pneumonia"

### Supplemental Materials

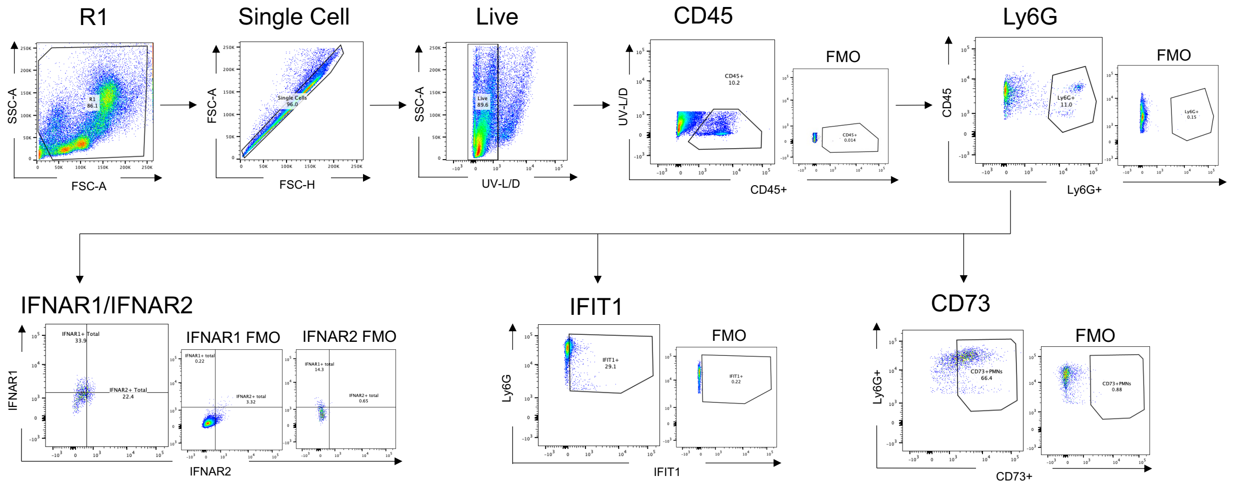


**Supplemental Figure 1. Gating Strategy for IFN Panel.**

Lungs and bone marrow from uninfected, +Sp (colonized alone), +IAV, and co-infected C57BL/6 (WT) or CD73KO mice were harvested and assessed for PMN phenoype by flow cytometry. Live single cells were gated on and the expression of IFNAR1, IFNAR2, IFIT1 or CD73 determined on PMNs (CD45+, Ly6G+). Fluorescent minus ones (FMOs) are depicted next to the representative flow plots for the indicated marker. Abbreviations: SSC-A = side scatter area; FSC-A = forward scatter area; FSC-H = forward scatter height; L/D=live-dead.

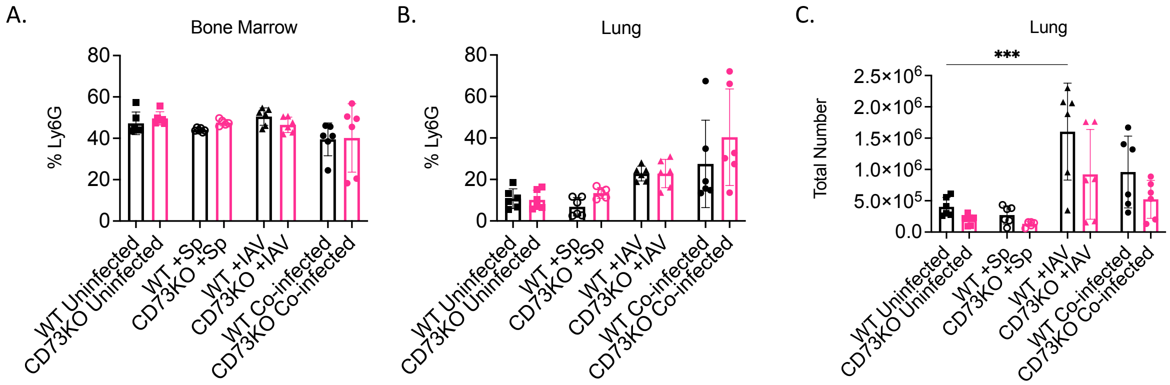


**Supplemental Figure 2. No difference in PMN presence in the bone marrow or influx to the lungs in response to infection in wild type versus CD73KO mice.**

WT and CD73KO mice were either co-infected with *S. pneumoniae* and Influenza A Virus (co-infected), infected with virus alone (+IAV), colonized with *S. pneumoniae* alone (+Sp), or mock-infected with PBS (uninfected). Two days post viral infection, the bone marrow and lungs were harvested and assessed by flow cytometry for percent and number of PMNs. The percent of PMNs in the bone marrow (A) and lungs (B), and total number of PMNs in the lungs (C) are shown. Data are pooled from two separate experiments with n=6 mice per group. *Denotes significant differences between indicated groups as determined by One-way ANOVA followed by Šídák's multiple comparisons test.

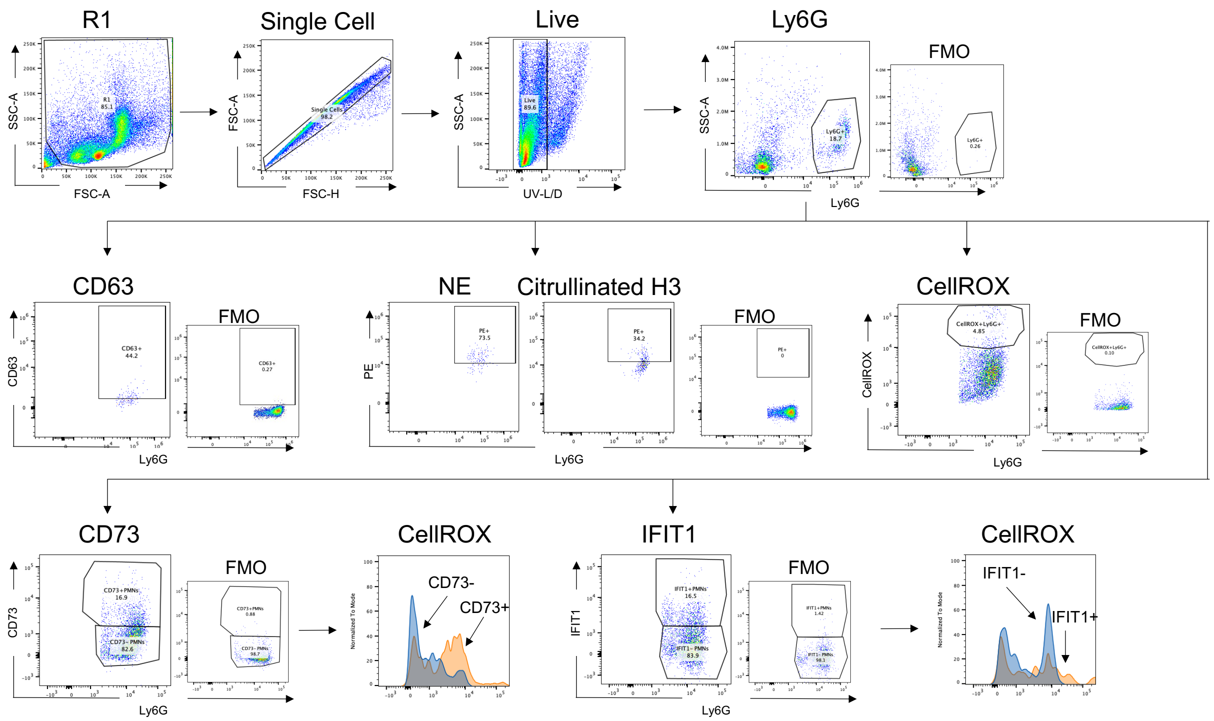


**Supplemental Figure 3. Gating strategy for PMN effector functions.**

Lungs and bone marrow from uninfected, +IAV, and co-infected WT or CD73KO mice were harvested and assessed for PMN phenoype by flow cytometry. Singlets were gated on and the expression of CellROX, CD73, IGIT1, CD63, neutrophil elastase (NE) or Citrullinated H3 (CitH3) determined on total PMNs (Ly6G+). The expression of CellROX under CD73+ versus CD73- and IFIT1+ versus IFIT- PMNs was also determined. Fluorescent minus ones (FMOs) are depicted next to the representative flow plots for the indicated marker. Abbreviations: SSC-A = side scatter area; FSC-A = forward scatter area; FSC-H = forward scatter height.

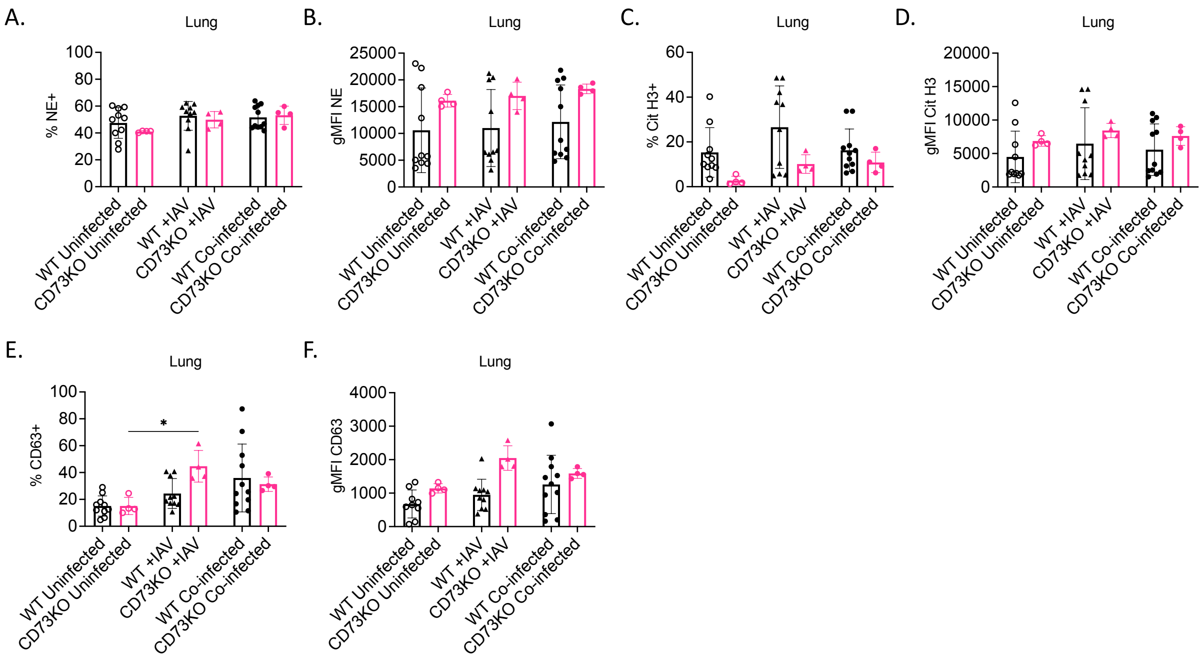


**Supplemental Figure 4. Pulmonary PMN effector activities in wild type versus CD73KO mice in response to *S. pneumoniae*/IAV co-infection.**

WT or CD73KO mice were either co-infected with *S. pneumoniae* and Influenza A Virus (co-infected), infected with virus alone (+IAV), or mock-infected with PBS (uninfected). Two days post IAV infection, the lungs were harvested and assessed by flow cytometry for percent % and expression (gMFI) of Neutrophil Elastase (NE) (A-B), Citrullinated H3 (CitH3) (C-D) and CD63 (E-F) on PMNs. Data are pooled from two separate experiments with n=4-8 mice per group. *Denotes significant differences as determined by Kruskal-Wallis test followed by Dunn's multiple comparisons test.

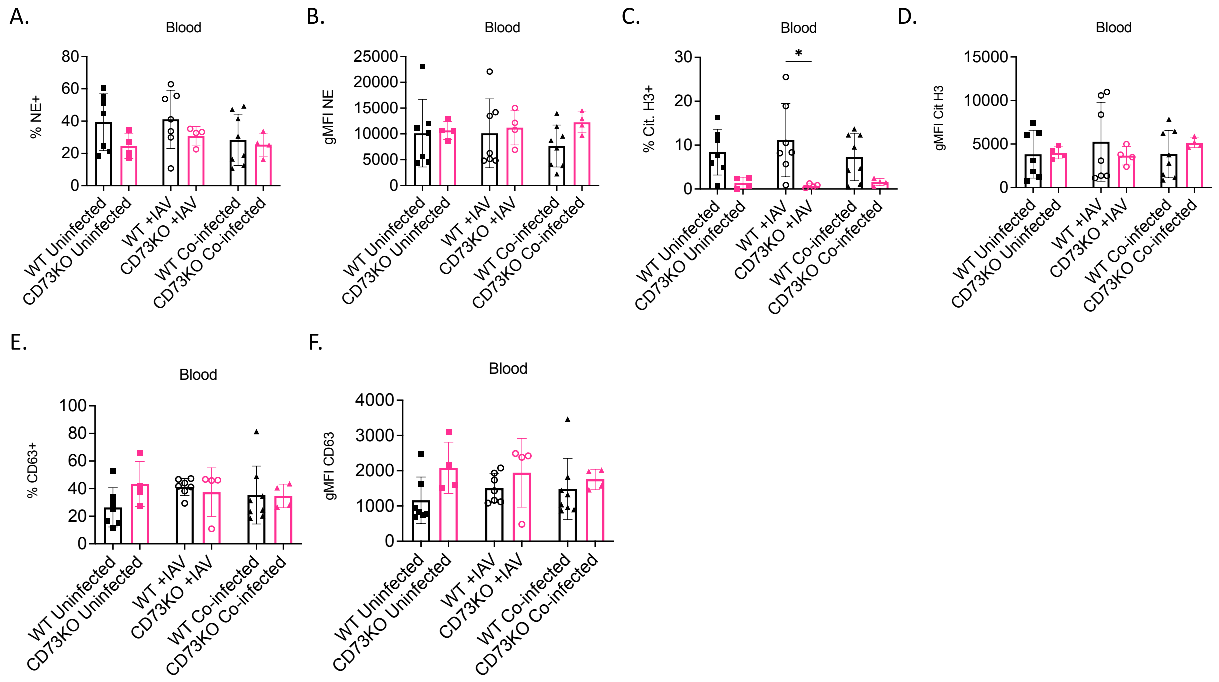


**Supplemental Figure 5. Circulating PMN effector activities in wild type versus CD73KO mice in response to *S. pneumoniae*/IAV co-infection.**

WT or CD73KO mice were either co-infected with *S. pneumoniae* and Influenza A Virus (co-infected), infected with virus alone (+IAV), or mock-infected with PBS (uninfected). Two days post IAV infection blood was harvested and assessed by flow cytometry for percent % and expression (gMFI) of Neutrophil Elastase (NE) (A-B), Citrullinated H3 (CitH3) (C-D) and CD63 (E-F) on PMNs. Data are pooled from two separate experiments with n=4-8 mice per group. *Denotes significant differences as determined by One-way ANOVA frollowed by Šídák’s multiple comparisons test.

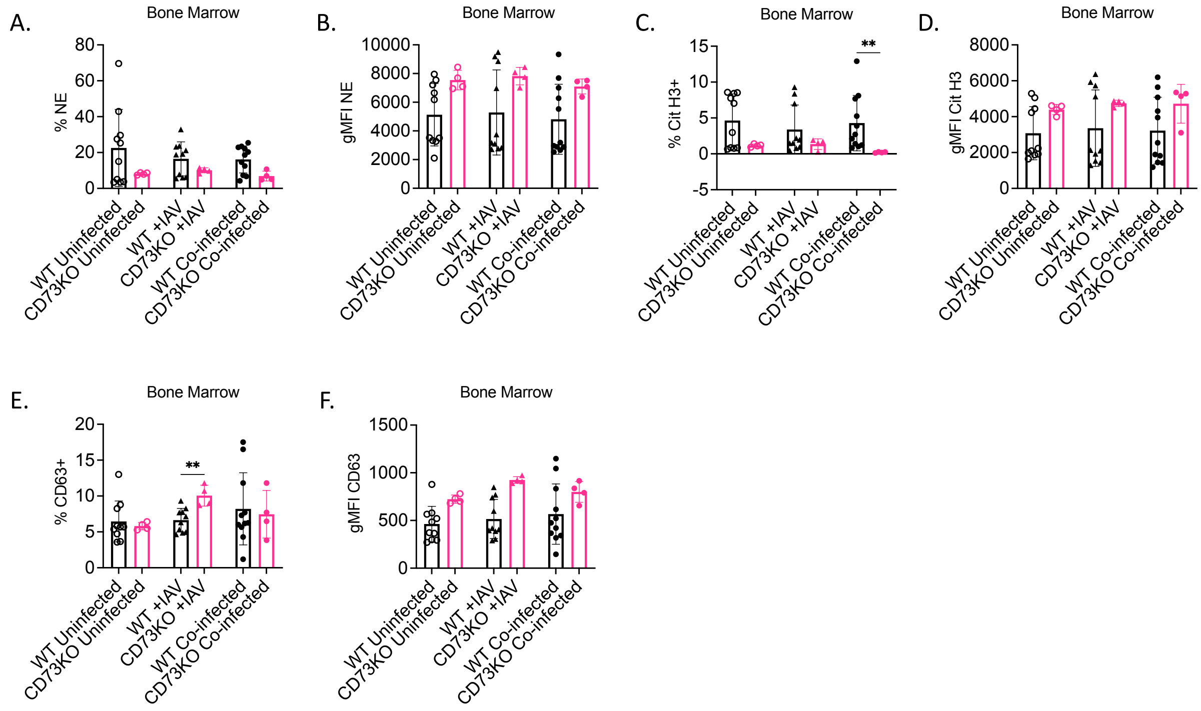


**Supplemental Figure 6. Bone marrow PMN effector activities in wild type versus CD73KO mice in response to *S. pneumoniae*/IAV co-infection.**

WT or CD73KO mice were either co-infected with *S. pneumoniae* and Influenza A Virus (co-infected), infected with virus alone (+IAV), or mock-infected with PBS (uninfected). Two days post IAV infection, the bone marrow was harvested and assessed by flow cytometry for percent % and expression (gMFI) of Neutrophil Elastase (NE) (A-B), Citrullinated H3 (CitH3) (C-D) and CD63 (E-F) on PMNs. Data are pooled from two separate experiments with n=4-8 mice per group. *Denotes significant differences as determined by One-way ANOVA frollowed by Šídák’s multiple comparisons test.
